# Biological insights from multi-omic analysis of 31 genomic risk loci for adult hearing difficulty

**DOI:** 10.1101/562405

**Authors:** Gurmannat Kalra, Beatrice Milon, Alex M. Casella, Yang Song, Brian R. Herb, Kevin Rose, Ronna Hertzano, Seth A. Ament

**Author notes:** <u>Lead Contact</u>: Seth A. Ament, Assistant Professor, Institute for Genome Science, HSF-3, 3^rd^ Floor, 670 W Baltimore St, Baltimore, MD 21201, Ronna Hertzano, Associate Professor, Department of Otorhinolaryngology-Head & Neck Surgery, 16 S Eutaw St. suite 500 – office, 800 West Baltimore St. Room 405 – laboratory.

## Abstract

Age-related hearing impairment (ARHI), one of the most common medical conditions, is strongly heritable, yet its genetic causes remain largely unknown. We conducted a meta-analysis of GWAS summary statistics from multiple hearing-related traits in the UK Biobank (n = up to 323,978) and identified 31 genome-wide significant risk loci for self-reported hearing difficulty (p < 5e-8), of which 30 have not been reported previously in the peer-reviewed literature at genome-wide significance. We investigated the regulatory and cell specific expression for these loci by generating mRNA-seq, ATAC-seq, and single-cell RNA-seq from cells in the mouse cochlea. Risk-associated genes were most strongly enriched for expression in cochlear epithelial cells, as well as for genes related to sensory perception and known Mendelian deafness genes, supporting their relevance to auditory function. Regions of the human genome homologous to open chromatin in sensory epithelial cells from the mouse were strongly enriched for heritable risk for hearing difficulty, even after adjusting for baseline effects of evolutionary conservation and cell-type nonspecific regulatory regions. Epigenomic and statistical fine-mapping most strongly supported 50 putative risk genes. Of these, at least 39 were expressed robustly in mouse cochlea and 16 were enriched specifically in sensory hair cells. These results reveal new risk loci and risk genes for hearing difficulty and suggest an important role for altered gene regulation in the cochlear sensory epithelium.

## INTRODUCTION

Age-related hearing impairment (ARHI) is characterized by a decline of auditory function due to impairments in the cochlear transduction of sound signals and affects approximately 25% of those aged 65-74 and 50% aged 75 and older^1^. Causes of ARHI and related forms of adult-onset hearing difficulty include a complex interplay between cochlear aging, noise exposure, genetic predisposition, and other health co-morbidities. Anatomical and physiological evidence suggest that these forms of hearing difficulty arise most commonly from damage to cochlear sensory epithelial cells, particularly inner and outer hair cells. Some forms of hearing difficulty also arise from damage to non-epithelial cells in the cochlea, including spiral ganglion neurons and cells of the stria vascularis.

Twin and family studies suggest that 25-75% of risk for ARHI is due to heritable causes^2^. Mutations in >100 genes cause monogenic deafness or hearing loss disorders^3^. However, a substantial fraction of patients with ARHI do not have a mutation in any known deafness gene, suggesting that additional genetic causes remain to be discovered.

Common genetic variation may contribute to these unexplained cases of hearing difficulty, but specific risk variants remain poorly characterized. Genome-wide association studies (GWAS) of hearing-related traits, including ARHI, tinnitus, and increased hearing thresholds, have identified only four genome-wide significant risk loci^4–8^. Positional candidate genes at these previously reported risk loci include *TRIOBP*, a gene associated with prelingual nonsyndromic hearing loss^4^; *ISG20*, encoding a protein involved in interferon signaling^4^; *PCDH20*, a member of the cadherin family^5^; and *SLC28A3*, a nucleoside transporter^5^. In addition, several studies have reported suggestive associations near *GRM7*, encoding a metabotropic glutamate receptor^6,8^.

Here, we performed multi-trait analyses of publicly available GWAS summary statistics from hearing-related traits in the UK Biobank (n up to 323,978) and identified 31 risk loci for hearing difficulty, of which 30 have not been described in peer-reviewed publications. We further characterized these risk loci using functional genomics data from the cochlea, including newly generated mRNA-seq, ATAC-seq, and single-cell RNA-seq from cochlear epithelial and non-epithelial cells of neonatal mice. Our results indicate that heritable risk for hearing difficulty is enriched in genes and putative enhancers expressed in sensory epithelial cells, as well as for common variants near Mendelian hearing loss genes. Statistical and epigenomic fine-mapping most strongly supported 50 putative risk genes at these loci, predicting both protein-coding and gene regulatory mechanisms for ARHI.

## RESULTS

### Heritability of hearing-related traits in the UK Biobank

The UK Biobank is a population-based, prospective study with over 500,000 participants in Britain, aged 40–69 years when recruited in 2006–2010^9^. The study has collected data on thousands of phenotypes, as well as genome-wide genotyping data. Recently, the Neale lab at Massachusetts General Hospital performed GWAS of up to 337,000 individuals and 2,419 traits in the UK Biobank and made the summary statistics and heritability estimates publicly available (http://www.nealelab.is/uk-biobank/). Using these data, we examined heritability of 31 hearing-related traits, including 14 self-reported traits and 17 traits derived from International Statistical Classification of Diseases and Related Health Problems (ICD-10) codes (Fig. 1a; Table S1). Four traits, all self-reported, had statistically significant heritability (h^2^) explained by genotyped and imputed single nucleotide polymorphisms (SNPs), based on linkage disequilibrium (LD) score regression (LDSC^10^), after correction for multiple testing (raw p-values < 2.1e-5; alpha = 0.05). These traits were “hearing difficulty/problems: Yes” (henceforth, “hearing difficulty”), “hearing difficulty/problems with background noise” (henceforth, “background noise problems”), “hearing aid user”, and “tinnitus: yes, now most or all of the time” (henceforth “tinnitus”).

**Figure 1.**
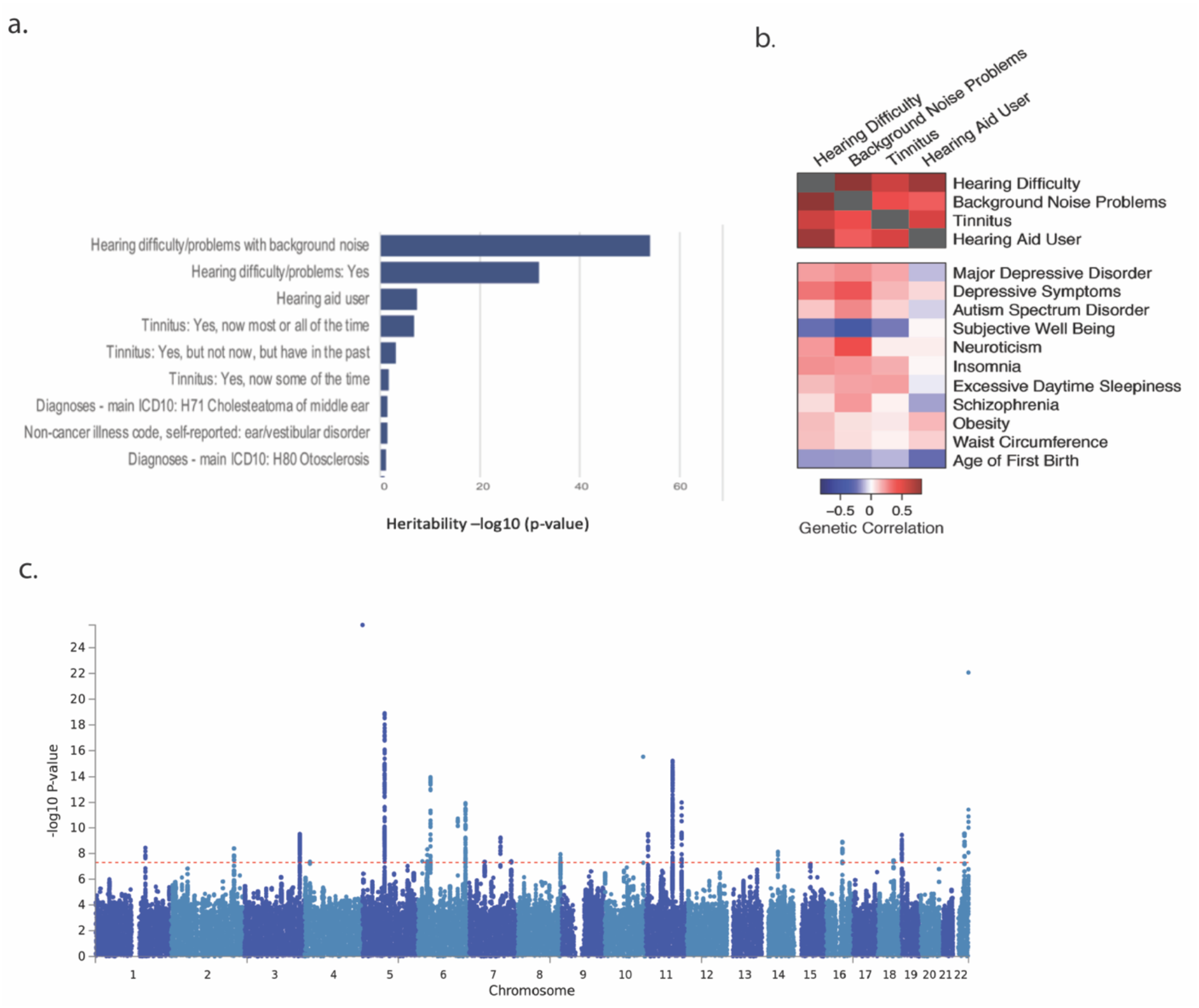
Genome-wide association studies of hearing-related traits in the UK Biobank. a. Heritability of top 9 hearing-related traits in the UK Biobank. b. Genetic correlations among the four most significantly heritable hearing-related traits and between these traits and 14 non-hearing traits. c. Manhattan plot for genetic associations with hearing difficulty in the UK Biobank, following meta-analysis across the four hearing-related traits.

We used LDSC to study the mean χ^2^ statistics for each trait, and we estimated the proportion of inflation due to polygenic heritability versus confounding. As expected, quantile-quantile plots indicate substantial deviation of χ^2^ statistics from a null distribution (Fig. S1). Background noise problems (Intercept = 1.031, Int. p = 1.3e-3) and hearing difficulty (Intercept = 1.018; Int. p = 3.7e-2) had significant LDSC intercept terms, suggesting some confounding, whereas the intercept terms for hearing aid use and tinnitus were not significant. Reassuringly, for all four traits, LDSC intercepts ascribe >90% of the inflation in the mean χ^2^ to polygenic heritability rather than to confounding. These results suggest that hearing-related traits in the UK Biobank are heritable and highly polygenic.

Next, we assessed whether risk for the four hearing-related traits arises from shared or distinct genetic factors. Using LDSC, we found that all pairs of hearing-related traits were genetically correlated (all r_g_ ≥ 0.37; all p-values < 7.1e-8; Table S2). Genetic correlations were strongest between the two most significantly heritable traits, hearing difficulty and background noise problems (r_g_ = 0.81). These results suggest substantial shared genetics among these hearing-related traits.

In addition, we assessed genetic correlations between the UKBB hearing-related traits and GWAS of 235 non-hearing traits, available via LD Hub^11^. As expected, genetic correlations among hearing traits were stronger than genetic correlations between hearing traits and non-hearing traits. In addition, we detected significant genetic correlations (False Discovery Rate < 5%) between hearing-related traits and 14 other traits (Fig. 1b, Table S2). Eleven of the 14 genetically correlated traits are psychiatric and personality traits, including positive genetic correlations of hearing difficulty with major depressive disorder and insomnia and a negative genetic correlation of hearing difficulty with subjective well-being. In addition, we detected positive genetic correlations between hearing difficulty and two anthropomorphic traits: obesity and waist circumference. Finally, we detected a negative genetic correlation between hearing difficulty and the age at first childbirth, a proxy for educational attainment and cognition. Genetic correlations typically arise from diverse direct and indirect relationships, yet, remarkably, many of these correlations reflect known comorbidities and risk factors for hearing loss^12,13^.

### Genomic risk loci

Leveraging the shared heritability among the four selected hearing-related traits, we performed a multi-trait analysis with MTAG (Multi-Trait Analysis of GWAS). MTAG uses GWAS summary statistics from multiple traits and can boost statistical power when the traits are genetically correlated^14^. The original GWAS summary statistics for hearing difficulty included genome-wide significant associations (p < 5e-8) of hearing difficulty with 779 SNPs at 22 approximately LD-independent genomic loci (Fig. 1c). Following joint analysis with MTAG, we identified genome-wide significant associations of hearing difficulty with 988 genotyped and imputed SNPs, located at 31 approximately LD-independent genomic loci (Table S3). In addition, MTAG analysis revealed 20 genome-wide significant loci for background noise problems, 25 for hearing aid use, and 20 for tinnitus (Tables S4-S6). Most of these loci overlap the 31 loci for hearing difficulty. In our subsequent analyses, we utilized MTAG summary statistics for hearing difficulty.

Next, we examined overlap between the 31 hearing difficulty risk loci in the UK Biobank sample versus previously reported GWAS of hearing-related traits. There were no suitably powered datasets available for replication of our findings, as the UK Biobank cohort is orders of magnitude larger than cohorts used in previously-reported GWAS. This issue is not unique to hearing-related traits, and it has become common to report findings from biobank-scale GWAS without standard replication^15,16^. Nonetheless, we considered whether findings from previous GWAS of hearing-related traits replicate in the UK Biobank. We analyzed 59 SNPs reported at genome-wide or suggestive significance levels in previous GWAS of hearing-related traits^4–8,17^ and which were genotyped or imputed in the UK Biobank sample. Eight of these 59 SNPs showed nominally significant associations with hearing difficulty in the UK Biobank (p < 0.05; Table S7), including both loci that reached genome-wide significance in the largest previous GWAS of ARHI^4^: rs4932196, 54 kb 3’ of ISG20 (p = 2.6e-5 in the UK Biobank); and rs5750477, in an intron of *TRIOBP* (p = 1.3e-6). Also replicated in our analysis were two SNPs previously reported at a suggestive significance level, in or near genes that cause Mendelian forms of hearing loss: rs9493627, a missense SNP in *EYA4*^4^ (p = 7.7e-10); rs2877561, a synonymous variant in *ILDR1*^4^ (p = 1.1e-8). In addition, we found a nominal level of support for rs11928865, in an intron of *GRM7*, previously reported at a suggestive significance level in multiple cohorts with ARHI^6,8^ (p = 2.2e-2). Taken together, these results suggest that hearing difficulty risk loci in the UK Biobank overlap with previous results, while vastly expanding our knowledge of risk loci related to ARHI.

### Heritability for hearing difficulty is enriched near Mendelian deafness genes and genes expressed in cochlear cell types

Next, we sought biological insights into hearing difficulty through gene set enrichment analyses. We performed gene-based analyses of the MTAG summary statistics using MAGMA^18^ (<u>M</u>ulti-marker <u>A</u>nalysis of <u>G</u>eno<u>M</u>ic <u>A</u>nnotation) and identified 104 genes reaching a genome-wide significance threshold, p < 2.5e-6, correcting for 20,000 tests (Table S8). Of these 104 genes, 40 overlap with the 31 risk loci, while the remaining genes are located at additional loci where no individual SNP reached genome-wide significance. We performed a series of hypothesis-based and exploratory gene set enrichment analyses.

It has been hypothesized that age-related hearing loss involves low penetrance variants in genes that are also associated with monogenic deafness disorders^4,19^. To test this hypothesis, we studied common-variant associations near 110 Mendelian deafness genes from the Online Mendelian Inheritance in Man (OMIM) database. These Mendelian deafness genes were enriched at hearing difficulty risk loci (p = 1.19e-6; Table S9). We detected gene-based associations with hearing difficulty at a nominal level of significance (p-values < 0.01) for 15 of these 110 genes, with the strongest associations at *TRIOBP* (MAGMA: p = 1.2e-10), *ILDR1* (p = 2.5e-8), and *MYO7A* (p = 8.5e-5). These findings support the hypothesis that Mendelian hearing loss genes contribute to age-related hearing difficulty, but also suggest that many risk loci for hearing difficulty involve genes that have not previously been implicated in hearing loss.

A more general hypothesis is that hearing difficulty risk is enriched in genes expressed in the cochlea. We generated mRNA-seq from FACS-sorted cochlear epithelial cells, cochlear mesenchymal cells, cochlear neurons, and cochlear vascular endothelial cells in the mouse cochlea. We calculated the median expression of each gene in each of these cell types, as well as in four subtypes of sensory epithelial hair cells and supporting cells derived from published RNA-seq^20–22^. For comparison, we considered the expression of each gene in 53 extracochlear human tissues and cell types from the Genotype-Tissue Expression consortium (GTEx)^23^. Using MAGMA gene property analysis, we tested for associations of tissue-specific expression levels with genetic risk for hearing difficulty. Risk for hearing difficulty was enriched in genes expressed in cochlear cell types, with the greatest enrichment in cochlear epithelial cells in general, followed by cochlear hair cells and cochlear non-epithelial cell types (Fig. 2; Tables S9,S10). The enrichments in cochlear epithelial cells are stronger than for any of the non-cochlear tissues. Therefore, our results suggest that many of the risk loci are explained by genes that are expressed in the cochlea.

**Figure 2.**
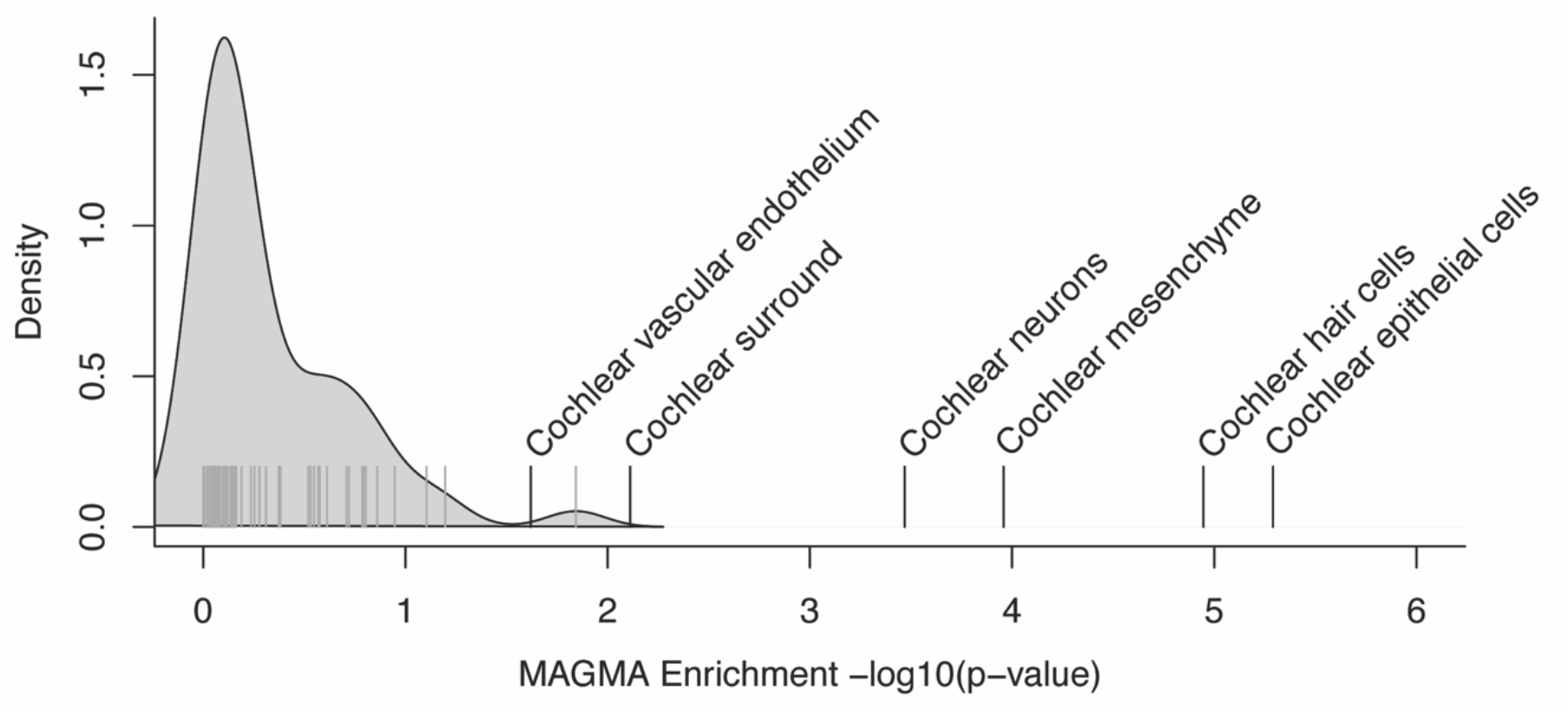
Heritable risk for hearing difficulty is enriched near genes expressed in the cochlea. Black vertical lines indicate the −log10(p-value) for the enrichment of hearing difficulty risk near genes expressed in each cochlear cell type. Gray vertical lines indicate −log10(p-value) for genes expressed in each of 53 non-cochlear tissues and cell types.

To identify additional functional categories enriched for hearing difficulty risk, we performed an exploratory analysis of 5,917 gene sets from Gene Ontology (GO). This analysis revealed a single significant GO term after correction for multiple testing: sensory perception of mechanical stimulus (150 genes in this set; p = 8.62e-9) (Table S11). Taken together, these results support the relevance of hearing difficulty risk loci to the auditory system, including many genes that have not previously been associated with hearing loss.

### Heritable risk for hearing difficulty is enriched in open chromatin regions from cochlear epithelial cells

Many studies have demonstrated that GWAS associations are enriched in gene regulatory regions such as enhancers and promoters, especially in regulatory elements that are active in disease-relevant tissues and cell types. For instance, SNPs associated with psychiatric disorders are enriched in regulatory regions active in the brain, and SNPs associated with autoimmune disorders are enriched in gene regulatory regions active in myeloid cells^24,25^. Consequently, we hypothesized that SNPs influencing risk for hearing difficulty are enriched in gene regulatory regions active in the cochlea.

Mice have previously been used to successfully identify new deafness related genes and inner ear development^26^. To identify gene regulatory regions in the cochlea, we FACS-sorted epithelial cells (CD326+; including hair cells and supporting cells) and non-epithelial cells (CD326−; predominantly mesenchymal cells) from mouse cochlea at postnatal day 2 (Fig. 3a), and performed ATAC-seq, on biological duplicates, to identify open chromatin regions in each cell type. We identified 228,781 open chromatin regions in epithelial cells and 433,516 in non-epithelial cells. 113,733 regions were unique to epithelial cells (2.83% of the mouse genome), 320,871 unique to non-epithelial cells (4.47% of the mouse genome), and 120,919 overlapping (Fig. 3b; Tables S12,S13). We validated these open chromatin regions through comparison to experimentally validated enhancers from the VISTA Enhancer Database with activity in the ear^27^. ATAC-sensitive regions from both epithelial and non-epithelial cells overlapped significantly with the 15 known ear enhancers from the Enhancer Browser Database (epithelial cells: 3.1-fold enriched, p < 1.0×10^−4^; non-epithelial cells: 2.9-fold enriched, p < 1.0×10^−4^ based on 10,000 permutations). Examination of known cell type-specific genes suggested that chromatin accessibility in epithelial versus non-epithelial cells was correlated with cell type-specific gene expression (Fig. 3c-e). For instance, we detected open chromatin specific to epithelial cells near *Epcam* and *Sox2*, which are expressed specifically in cochlear epithelial cells^28,29^; and open chromatin specific to non-epithelial cells around *Pou3f4*, a marker for non-epithelial cells^30^.

**Figure 3.**
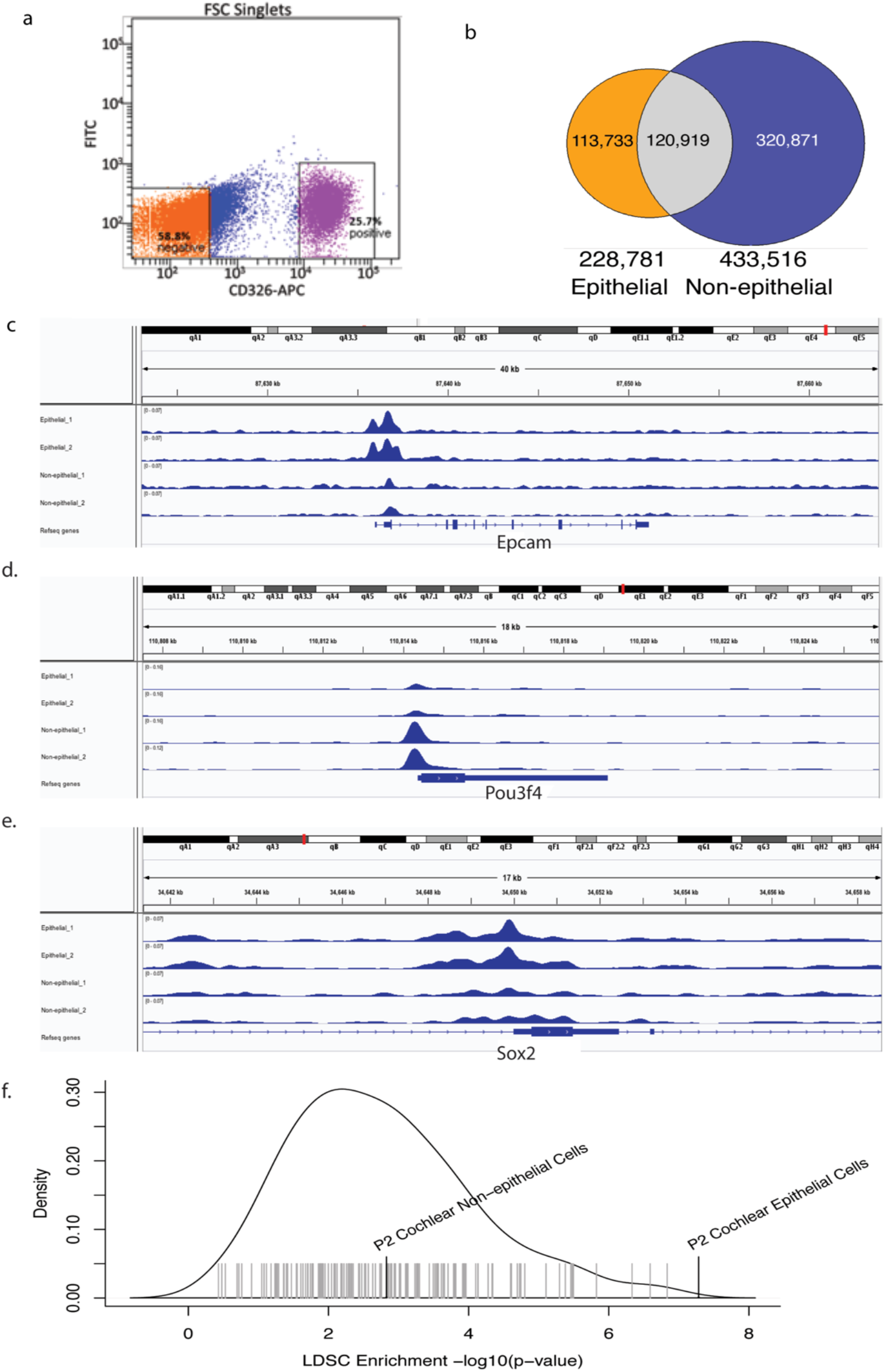
Heritable risk for hearing difficulty is enriched at open chromatin regions in cochlear sensory epithelial cells. a. FACS sorting of cochlear cells. Cochlear cells were labeled with a CD326 antibody conjugated to APC, and sorted two ways as CD326 (+) and CD326 (−). b. Overlap of open chromatin regions identified by ATAC-seq of sensory epithelial vs. non-epithelial cells in the mouse cochlea. c-e. Open chromatin peaks near cell type-specific marker genes: *Epcam* (b), *Pou3f4* (b), and *Sox2* (c). f. −log10(p-value) for enrichment of hearing difficulty risk in regions of the human genome homologous to open chromatin in epithelial and non-epithelial cells from mouse cochlea (black vertical lines) and in non-cochlear cell types from ENCODE (gray lines).

Next, we asked whether these putative regulatory regions in the cochlea are enriched for SNPs associated with hearing difficulty. Using the UCSC LiftOver tool^31^, we mapped ATAC-sensitive regions from each cochlear tissue type to the human genome to identify homologous genomic regions. 55.5% of the mouse epithelial regions and 50.2% of non-epithelial regions mapped to the human genome. We tested for enrichment of hearing difficulty risk in these conserved cochlear regions using stratified LD score regression^32^. This model tests for heritability in cochlea-specific regions after accounting for a baseline model consisting of 24 non-cell type-specific genomic annotations, including evolutionarily conserved regions and regions that are open chromatin across many tissues. Heritable risk for hearing difficulty was enriched 9-fold in epithelial open chromatin regions (Fig. 3f). 2.1% of all SNPs are in the annotated regions, and these SNPs capture 19.5% of the total SNP heritability (p = 5.2e-8). Heritability was less strongly enriched in open chromatin regions from non-epithelial cells (4.6-fold enriched; 3.0% of all SNPs are in the annotated regions, and the SNPs capture 14.2% of the total heritability; p = 0.001). For comparison, we performed comparable analyses using open chromatin regions from 147 DNase-seq experiments in 42 mouse tissues and cell types, generated by the ENCODE project^33^ (Table S14). The significance of the heritability enrichment in cochlear epithelial cells was greater than for any of the non-cochlear tissues. These results suggest that heritable risk for hearing difficulty is enriched specifically in evolutionarily conserved gene regulatory regions active in cochlear epithelial cells.

### Statistical and epigenomic fine-mapping supports functional consequences to 50 genes at hearing difficulty risk loci

Next, we sought to predict causal variants and target genes at each of the 31 hearing difficulty risk loci, considering both protein-coding and putative gene regulatory consequences of each variant. To begin this analysis, we identified 613 SNPs and short indels that are in strong LD (r^2^ > 0.9) with a genome-wide significant lead SNP at one of the 31 risk loci (Table S15).

We scanned these 613 risk variants for non-synonymous and stopgain SNPs, frameshift and non-frameshift indels, and effects on splice donor and acceptor sites, focusing on those variants that are predicted to be deleterious with a CADD Phred score > 10. We identified nine such protein-coding variants, including missense SNPs in *CLRN2, CRIP3, EYA4, CHMP4C, TYR, TRIOBP (2x), BAIAP2L2*, and *KLHDC7B* (Table 1). Notably, six of these nine SNPs are LD-independent lead SNPs at their respective loci, increasing the statistical likelihood that these variants are causal for hearing difficulty risk. The missense SNPs in *TRIOBP* and *BAIAP2L2* are annotated to the same risk locus at 22q13.1 (Fig. S3). The two *TRIOBP* variants are in strong LD (r^2^ = 0.97), so their effects may be additive or synergistic. By contrast, the *BAIAP2L2* variant is not in LD with either of the *TRIOBP* variants, suggesting an independent effect.

**Table 1.**
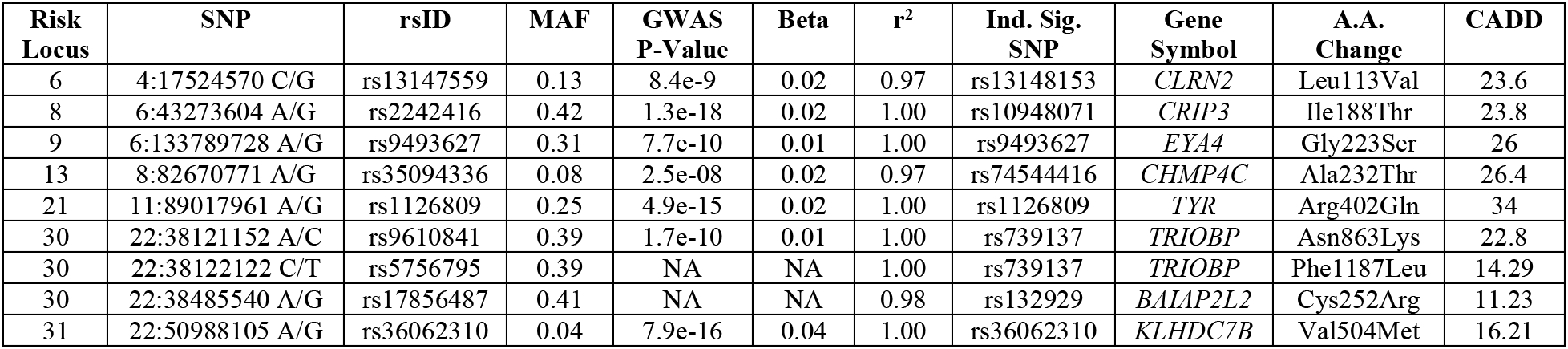
Deleterious protein-coding variants in strong LD with LD-independent genome-wide significant SNPs associated with hearing difficulty.

Causal variants on risk haplotypes that do not contain protein-coding variants may alter gene regulation. To elucidate these gene regulatory consequences, we annotated the 613 risk-associated SNPs in the context of the local two-dimensional and three-dimensional chromatin architecture. For the former, we utilized our ATAC-seq data from cochlear cells. For the latter, we used publicly available Hi-C data from 20 non-cochlear tissues and cell types^34^. Since data for chromatin architecture in the cochlea is unavailable, we considered all cell types in aggregate and set a stringent chromatin interaction significance threshold (p-value < 1e-25) to focus on the strongest chromatin loops. 126 of the 613 SNPs had potential gene regulatory functions in cochlea, based on homology to open chromatin in cochlear epithelial and non-epithelial cells from neonatal mice (Table S15). 57 of these 126 SNPs were located proximal (<10 kb) to the transcription start sites of 17 potential target genes. In addition, 100 of the 126 SNPs could be assigned to 72 distal target genes based on long-distance chromatin loops that connect the regions containing risk-associated SNPs to these genes’ transcription start sites located up to 3 Mb away (Table S16).

We integrated the coding and non-coding functional annotations to prioritize the most likely causal genes at each locus. The union of functional annotations supported 84 genes (Table S17). We prioritized 50 of these genes, as follows: (i) if one or more genes at a locus contained risk-associated protein-coding variants, we selected those genes; (ii) if no coding variants were identified at a locus, we selected the proximal target gene(s) of non-coding SNPs with predicted regulatory functions; (iii) if no proximal genes were identified, we considered distal target genes.

This analysis identified putative risk genes at 19 of the 31 risk loci, including 10 loci at which a single gene appears most likely to be causal (Table S18).

We sought independent support for roles of these genes in hearing loss or cochlear function based on prior evidence from genetic studies in humans and mice. Rare mutations in five of the 50 genes have been shown previously to cause Mendelian forms of deafness or hearing loss: *TRIOBP, EYA4, FTO, SOX2*, and LMX1A^4,35–43^. Genetic studies in mice have demonstrated hearing loss or cochlear development phenotypes for an additional seven of the 50 genes: *SYNJ2, TYR, PTGDR, MMP2, RPGRIP1L, RBL2*, and BAIAPL2^44–50^. Notably, three of the five genes with independent support from human rare variants -- *FTO, SOX2* (Fig. 4a), and *LMX1A* (Fig. 4b) -- are located distal to risk-associated SNPs and were predicted as target genes based on long-distance chromatin interactions, validating this approach for predicting causal mechanisms. We note that there may be additional causal genes for which the functional variants are missed by our analysis. For instance, at the 3q13.3 risk locus our approach excludes a strong positional candidate, *ILDR1*, in which loss-of-function variants cause a recessive hearing loss disorder^37^, since none of the risk variants at this locus were predicted to alter *ILDR1* function.

**Figure 4.**
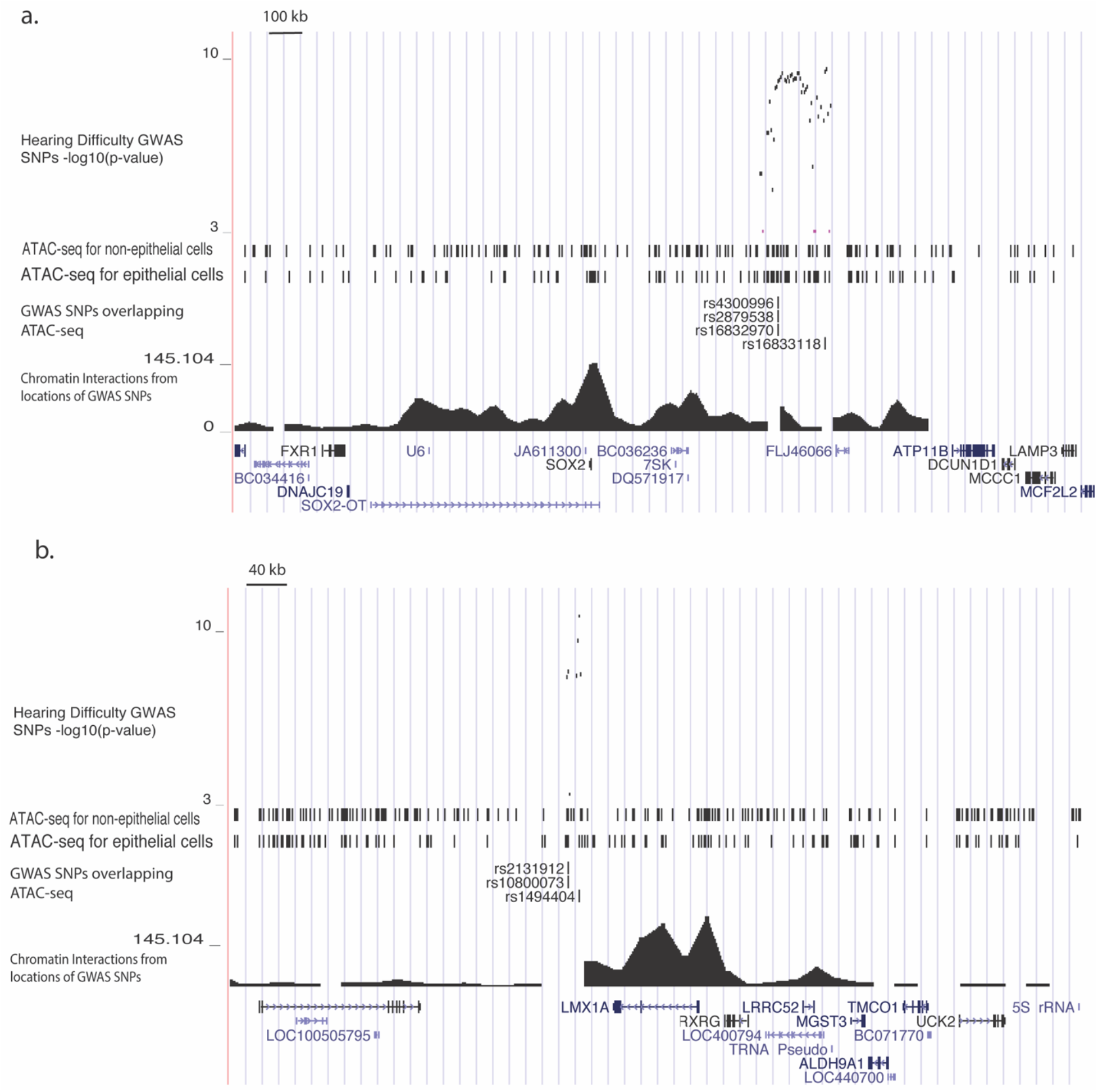
Epigenomic fine-mapping predicts distal target genes for hearing difficulty risk loci. Genetic associations and epigenomic annotations at chr3q26.3 (a) and chr1q23.3 (b). From top to bottom, genome browser tracks indicate: −log10(p-values) for association with hearing difficulty; locations of putative open chromatin in cochlear epithelial and non-epithelial cells, based on homology to ATAC-seq peaks in these cell types from neonatal mice; fine-mapped SNPs in strong LD with an LD-independent lead SNP and located <500bp from a putative cochlear open chromatin region; −log10(p-values) for chromatin interactions between the locations of the fine-mapped SNPs and distal regions, based on the minimum chromatin interaction p-value in each 40kb region from Hi-C of 20 non-cochlear human tissues and cell types; locations of UCSC knownGene gene models.

### Hearing difficulty risk genes are expressed in diverse cochlear cell types

To better understand the potential functions of the 50 putative risk genes in the cochlea, we investigated their expression patterns in cochlear cell types. We sequenced the transcriptomes of 3,411 single-cells from the mouse cochlea (postnatal day 2) using 10x Genomics Chromium technology. Cells were sequenced to a mean depth of 107,590 reads, which mapped to a median of 1,986 genes per cell. After quality control (Methods), we analyzed data from 3,314 cells. Louvain modularity clustering implemented with Seurat^51^ revealed 12 major clusters of cells (Fig 5a, Fig S2). Based on the expression of known marker genes (Table S19), we assigned these cell clusters to the following cell types: three clusters of epithelial cells (*Epcam*+; n = 419, 101, and 24 cells per cluster), three clusters of mesenchymal cells (*Pou3f4*+; n = 887, 701, and 76 cells per cluster, of which the smallest cluster are *2810417H13Rik*+ cells undergoing cell division), 324 glial cells (*Mbp*+), 391 medial interdental cells (*Otoa*+), 59 inner ear progenitors (*Oc90*+), 79 vascular cells (*Cd34*+), 161 sensory epithelial supporting cells (*Sox2*+), and 91 sensory hair cells (*Pou4f3*+).

**Figure 5.**
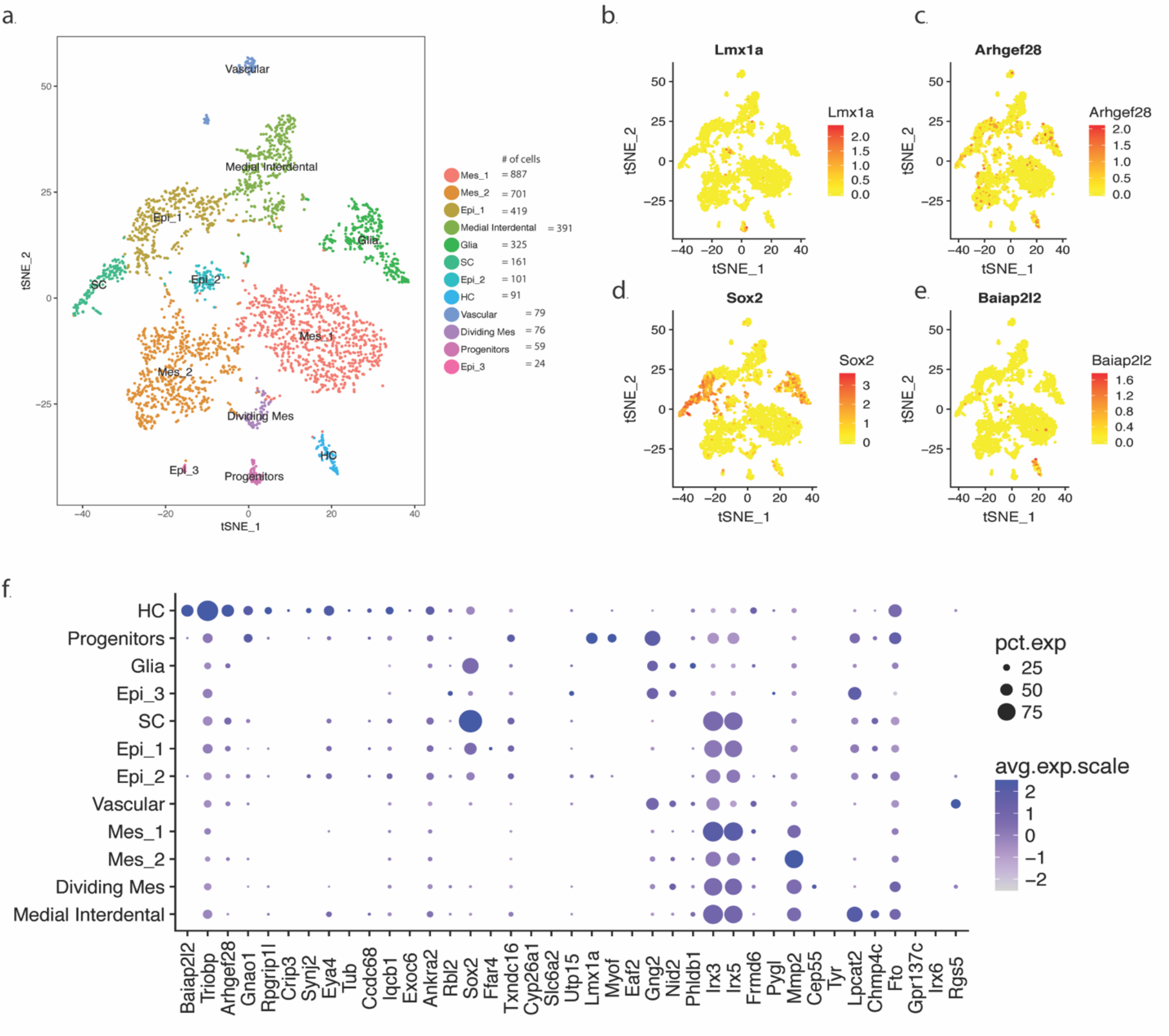
Single-cell RNA-seq of mouse cochlea reveals cell type-specific expression patterns of hearing difficulty risk genes. a. t-distributed stochastic neighbor embedding (t-SNE) plot of 3,411 cells in the postnatal day 2 mouse cochlea colored by Louvain modularity clusters corresponding to 12 cell types. b-e. t-SNE plots colored by the expression of selected hearing difficulty risk genes expressed selectively in cochlear cell types: *LMX1A* in sensory progenitor cells (b); *ARHGEF28* in hair cells (c), *SOX2* in supporting cells (d), and *BAIAP2L2* in hair cells (e). f. Dot plot showing the average expression and percent of cells with non-zero counts for each cochlea-expressed risk gene in each of the 12 cochlear cell types.

We tested cell type specific expression for each hearing difficulty risk gene. 39 of the 50 risk genes were expressed highly enough in these cochlear cells to be included in this analysis. By far the largest number of genes, 14 out of 39, were expressed selectively in sensory hair cells (Fig. 5f; Table S20). These hair cell-specific risk genes included known hearing loss genes such as *Triobp* (p = 9.5e-42) and *Eya4* (p = 2.3e-21), as well as genes that have not previously been implicated in human hearing loss; e.g., *Baiap2l2* (p = 2.6e-192; Fig. 5e), *Arhgef28* (p = 9.8e-31; Fig. 5c), *Gnao1* (p = 4.1e-29), *Rpgrip1l* (p = 6.3e-24), and *Crip3* (p = 9.1e-21). We also found risk genes that were expressed selectively in other cell types, including sensory epithelium supporting cells (*Sox2*, p = 9.5e=151; Fig. 5d), sensory epithelium progenitor cells (e.g., *Lmx1a*, p = 7.4e-131; Fig. 5b), medial interdental cells (*Lpcat2*, p = 1.8e-139), and mesenchymal cells (*Mmp2*, p = 6.0e-120).

We sought to corroborate the expression pattern of risk genes in hair cells using published expression profiles from hair cells isolated by three other methods: (i) transcriptome profiling of FACS-purified hair cells versus surrounding cells^52^; (ii) translatome profiling of RiboTag-purified hair cells versus surrounding cells^53^; and (iii) single-cell RNA-seq of sensory epithelial cells^54^. Meta-analysis of these three datasets confirmed selective expression in hair cells (FDR < 0.1) for 12 of the 14 genes above (Table S21). This analysis also revealed low but highly specific expression in hair cells for two other risk genes, *Clrn2* and *Klhdc7b*. Thus, in total, we find that 16 of the 50 putative hearing difficulty risk genes identified by GWAS are expressed selectively in sensory hair cells.

## DISCUSSION

Here, we have described a well-powered GWAS of hearing difficulty, leveraging results from >300,000 participants in the UK Biobank, and we interpreted these genetic associations in the context of multi-omic data from the mouse cochlea. We identified 31 risk loci for hearing difficulty, of which 30 had not reached genome-wide significance in any previous GWAS of hearing-related traits. Heritable risk for hearing difficulty was enriched in genes and gene regulatory regions expressed in sensory epithelial cells, as well as for common variants near Mendelian hearing loss genes. We identified 50 putative risk genes at these loci, many of which were expressed selectively in sensory hair cells and other disease-relevant cell types.

Prior to this work, a prominent hypothesis had been that age-related hearing impairment involves lower-penetrance genetic variation in genes that cause Mendelian forms of hearing loss^4,19^. In support of this view, we found that heritability for hearing difficulty was enriched near Mendelian hearing loss genes, including genome-wide significant risk loci that overlapped three Mendelian hearing loss genes, *TRIOBP, EYA4*, and *ILDR1*. However, these signals represent a small fraction of the heritable risk. Indeed, our results better support a highly polygenic genetic architecture for hearing difficulty, spanning many genes that had not been known to influence hearing. Evidence for polygenicity includes the inflation of χ^2^ statistics (i.e., low p-values) across many thousands of SNPs and the enrichment of heritability across thousands of genes and putative gene regulatory regions expressed in the cochlear sensory epithelium. It is likely that the 31 risk loci identified here represent merely the tip of a larger genetic iceberg. Thus, as with other common traits^55^, it is likely that additional risk loci and risk genes will be discovered as sample sizes for GWAS continue to grow larger.

Our results suggest that genetic risk factors for hearing difficulty act most frequently – but not exclusively -- through mechanisms within sensory hair cells. The primacy of hair cells is supported by heritability enrichments for genes expressed in hair cells and for regions of open chromatin in the cochlear sensory epithelium, as well as by the fact that 16 of the 50 fine-mapped risk genes were expressed selectively in hair cells. These results strongly support the relevance of our findings to auditory function, since damage to hair cells is the most common pathophysiology in AHRI. Of the 16 hair cell-specific risk genes identified in our analysis, loss-of-function mutations in two are known to cause Mendelian hearing loss: *TRIOBP*^4,42,43^ and *EYA4*^39,40^. An additional five genes have not previously been associated with human hearing loss but are known to cause hearing loss or cochlear development when mutated in mice: *SYNJ2*^44^, *RPGRIP1L*^48^, *BAIAP2L2*^50^, *TUB*^56^, and *RBL2*^49^. To our knowledge, this is the first report of a hearing loss phenotype for the remaining seven hair cell-specific risk genes: *ANKRA2, ARHGEF28, CRIP3, CCDC68, EXOC6, GNAO1, IQCB1*, and *KLHDC7B*. Hearing difficulty risk haplotypes contained protein-coding variants in 6 of these genes, while the others were supported by non-coding variants with predicted gene regulatory functions. These genes have diverse biological functions, ranging from transcriptional regulation to intracellular signaling to metabolic enzymes to structural components of synapses and stereocilia. Taken together, these results suggest that functional variants impacting a wide range of hair cell-specific genes contribute to risk for hearing loss.

Other risk genes suggest plausible mechanisms for hearing loss involving components of the stria vascularis. The lead SNP at a risk locus on chr11q14.3 is a deleterious missense SNP in *TYR. TYR* encodes tyrosinase, which catalyzes the production of melanin. In the cochlea, *TYR* is expressed specifically in melanin-producing intermediate cells of the stria vascularis, and *TYR* mutant mice have strial albinism, accompanied by age-associated marginal cell loss and endocochlear potential decline^45^. Chromosomal contacts at chr16q12.2 suggest that *MMP2* is targeted by two distinct risk loci. *MMP2* encodes matrix metalloproteinase-2, which serves an essential role in the cochlear response to acoustic trauma by regulating the functional integrity of the blood-labyrinth barrier^47^. Risk-associated variants in these genes may contribute to strial atrophy, a common non-sensory cause of age-related hearing impairment^57^.

Perhaps more surprisingly, several of the risk genes identified in our analysis are best known for their functions in cochlear development. These include two well-characterized transcription factors: *SOX2* and *LMX1A*. Loss-of-function mutations in each of these genes cause deformations of the cochlea and hearing loss in humans^36,38^, while our analyses of adult hearing difficulty revealed non-coding genetic variation in putative distal enhancers. *SOX2* is required for the formation of prosensory domains that give rise to hair cells and supporting cells^58^. *LMX1A* maintains proper neurogenic, sensory, and non-sensory domains in the mammalian inner ear, in part by restricting and sharpening *SOX2* expression^59^. In addition, two genes supported by putative gene regulatory interactions at the chr10q23.3 risk locus, *CYP26A1* and *CYP26C1*, metabolize retinoic acid and are involved in the specification of the otic anterior-posterior axis^60^. Changes in cochlear development may cause vulnerabilities to hearing loss later in life. Alternatively, there may be as yet undescribed roles for these genes in the adult cochlea contemporaneous with hearing loss.

In summary, we report a well-powered GWAS of hearing difficulty, as well as new transcriptomic and epigenomic data from the mouse cochlea, revealing 31 risk loci and 50 putative risk genes. Previous GWAS of hearing-related traits had revealed only four genome-wide significant risk loci. Functional studies of these risk genes are warranted, especially for several hair cell-specific genes that had not previously been implicated in hearing loss. In addition, our findings support a polygenic genetic architecture for hearing difficulty, suggesting that more risk genes will be discovered as genetic data become available from additional biobank-scale cohorts.

## ACKNOWLEDGEMENTS

This work was supported by two grants from the Hearing Restoration Project of the Hearing Health Foundation (S.A.A., PI, R.H., PI) and by the National Institute of Deafness and Other Communication Disorders (R01 DC013817, R.H., PI; R00 DC013107, Thomas Coate, PI). We are grateful to Benjamin Neale and the leaders of the UK Biobank for making the GWAS resource freely available to the research community, as well as to all the participants in the UK Biobank, without whom none of this work would be possible.

## AUTHOR CONTRIBUTIONS

G.K., R.H., and S.A.A. designed research. B.M. and R.H. performed experiments with mice and prepared samples for sequencing. G.K., Y.S., A.M.C., B.R.H., K.R., R.H., and S.A.A. analyzed the data. G.K. and S.A.A. wrote the manuscript. All authors edited the manuscript.

## METHODS

### Animals

All procedures involving animals were carried out in accordance with the National Institutes of Health Guide for the Care and Use of Laboratory Animals and were approved by the Animal Care Committee at the University of Maryland (protocol numbers 0915006 and 1015003).

### Cell sorting by FACS followed by mRNA-seq and ATAC-seq

CD-1 timed-pregnant females were purchased from Charles River (Maryland). At postnatal day 2, the mice were euthanized and their temporal bone removed. Cochlear ducts from 20 mice were harvested, pooled and processed for Fluorescence-activated cell sorting (FACS) as described^28^. Briefly, to generate cell population for mRNA-seq, single-cell suspensions were obtained and incubated with anti-CD326, anti-CD49f, and anti-CD34 antibodies to detect sensory epithelial, neuronal, mesenchymal, and vascular endothelial cells. For ATAC-seq, a simplified protocol was utilized to distinguish sensory epithelial from non-epithelial cells (primarily mesenchyme) based on labeling with anti-CD326. Cells were sorted by FACS using a BD FACS Aria II Cell Sorter (BD Biosciences) at the Flow Cytometry Facility, Center for Innovative Biomedical Resources (University of Maryland School of Medicine) (Fig. 3a).

For mRNA-seq, libraries derived from total RNA from sorted cells were sequenced on an Illumina sequencer at the Institute for Genome Sciences (IGS) of the University of Maryland, School of Medicine. For ATAC-seq, fifty thousand cells and one hundred thousand cells from each sample were further processed as described ^61^ with the following modification: following the transposition reaction and purification step, a right side size selection (ratio 0.6) using SPRIselect (Beckman-Coulter, Indiana) was added before proceeding to the PCR amplification. This extra step resulted in the selection of DNA fragments between 150 bp to 700 bp. The following primers from ^61^ were used for library preparations:

> Ad1_noMX 5’-AATGATACGGCGACCACCGAGATCTACACTCGTCGGCAGCGTCAGATGTG-3’; Ad2.1_TAAGGCGA 5’-CAAGCAGAAGACGGCATACGAGATTCGCCTTAGTCTCGTGGGCTCGGAGATGT-3’; Ad2.2_CGTACTAG 5’-CAAGCAGAAGACGGCATACGAGATCTAGTACGGTCTCGTGGGCTCGGAGATGT-3’; Ad2.3_AGGCAGAA 5’-CAAGCAGAAGACGGCATACGAGATTTCTGCCTGTCTCGTGGGCTCGGAGATGT-3’; Ad2.4_TCCTGAGC 5’-CAAGCAGAAGACGGCATACGAGATGCTCAGGAGTCTCGTGGGCTCGGAGATGT-3’.

After completion of the libraries, whole genome sequencing, paired-end and a depth of 66 million reads, was performed on an Illumina HiSeq 4000 at IGS.

### Single-cell RNA sequencing of mouse cochlea

At postnatal day 2, 3 pups from a CD-1 timed-pregnant female were euthanized and their temporal bone removed. Cochlear ducts were harvested and pooled into Thermolysin (Sigma-Aldrich) for 20 min at 37°C. The Thermolysin was then replaced with Accutase (Sigma-Aldrich) and the tissue incubated for 3 min at 37°C followed by mechanical dissociation, repeating this step 3 times. After inactivation of the Accutase with 5% fetal bovine serum, the cell suspension was filter through a 35μm nylon mesh to remove cell clumps. The cell suspension was then processed for single-cell RNAseq.

Droplet-based molecular barcoding and single-cell sequencing were performed at the Institute for Genome Sciences (IGS) of the University of Maryland, School of Medicine. Approximately 10,000 dissociated cochlear cells were loaded into a Chromium Controller (10x Genomics) for droplet-based molecular barcoding of RNA from single cells. A sequencing library produced using the 10x Single Cell Gene Expression Solution. Libraries from two cochlear samples were sequenced across three lanes of an Illumina HiSeq4000 sequencer to produce paired-end 75 bp reads.

### Genetic correlations of hearing-related and non-hearing-related traits

We began by examining publicly available LDSC heritability estimates for 31 manually-selected hearing-related traits from the UK Biobank, based on the September 20, 2017, data release from the Neale lab (http://www.nealelab.is/blog/2017/7/19/rapid-gwas-of-thousands-of-phenotypes-for-337000-samples-in-the-uk-biobank). Four of these traits had significant heritability and were selected for further analysis: hearing difficulty (2407_1), background noise problems (2257), hearing aid use (3393), and tinnitus most or all of the time (4803_11). Genetic correlations among the four hearing-related traits were calculated on the original GWAS summary statistics from the Neale lab using LDSC genetic correlation analysis with default parameters. Genetic correlations of hearing-related traits with 328 additional traits was performed using LDHub v1.9.0 (http://ldsc.broadinstitute.org/ldhub/)^11^.

### Meta-analysis of hearing-related traits in the UK Biobank

Meta-analysis of the four hearing-related traits was performed with Multi-Trait Analysis of GWAS (MTAG) ^14^ using default parameters. MTAG is explicitly designed for joint analysis of summary statistics from biobank-scale GWAS of genetically correlated traits in overlapping samples.

### Replication of genetic associations in independent cohorts

We tested for replication of previously reported risk loci for ARHI and other hearing-related traits through lookups in the MTAG hearing difficulty summary statistics. We started with 62 previously-reported SNPs, derived from top-level results reported by Hoffmann et al. (2016)^4^, Vuckovic et al. (2015)^5^, and from several earlier studies as reported in Table S1 from Ref. 4. Summary statistics for 59 of these 62 SNPs were available in the UK Biobank sample.

### Gene set enrichment analysis

Gene set enrichment analyses were performed using MAGMA^18^ v1.06 implemented within FUMA^62^ v1.3.1, as well as via a standalone installation. Genotyped and imputed SNPs were annotated to ENSEMBL v92 gene models in FUMA. Annotations were limited to protein-coding genes, excluding the major histocompatibility (MHC) region of extended linkage disequilibrium (a common source of false positive results), and with SNPs mapping to a gene if they were located between the gene’s start and end position. MAGMA was then used to calculate a p-value for each gene, based on the mean association among the SNPs annotated to each gene. Gene-based p-values were used to perform the following gene-level analyses: (i) gene set enrichment analyses with Gene Ontology terms; (ii) gene set enrichment analysis with Mendelian deafness genes extracted from the Online Mendelian Inheritance in Man (OMIM) database (https://omim.org/); (iii) gene property analyses to assess covariance of MAGMA gene p-values with gene expression in six cochlear cell types and 53 non-cochlear cell types. For cochlear cell types, we computed the median transcripts per million (TPM) expression level for each gene in each cell type in RNA-seq of FACS-sorted cells from GSE64543^63^ and GSE60019^20^. For non-cochlear cell types, we downloaded median TPM values for 53 tissues from the GTEx v7 portal (gtexportal.org). We identified the set of genes quantified in all datasets and performed a quantile normalization across log-transformed TPM values from all cell types. Using these normalized expression levels, we performed a one-sided MAGMA gene property analysis, conditioning on the median expression level of each gene across all cell types, as well as standard covariate due to gene length, the number of SNP annotated to each gene, and correlations among nearby genes due to LD.

### ATAC-seq data processing

Four ATAC-seq fastq files (two epithelial and 2 non-epithelial samples from P1 mouse cochlea) from each tissue type were aligned to mm10 genome using BWA aligner *bwa mem* method (https://github.com/lh3/bwa). Sorted BAM files from each of the four samples were filtered to mapped reads only using samtools, converted to BED format using bedtools, and analyzed for open chromatin signal enrichment using F-Seq^64^. The two BED files for each tissue type were merged using *bedtools intersect*, to identify regions common to both samples, requiring at least a 1 base pair overlap. We removed blacklist regions computed by the ENCODE project (ENCFF547MET), which show high non-specific signal across many assays.

### Determining enrichment of ATAC-seq peaks for known tissue specific enhancers

We examined overlap between open chromatin regions from ATAC-seq experiments and tissue-specific enhancers from VISTA (https://enhancer.lbl.gov/) using the Genomic Association Tester (GAT; https://github.com/AndreasHeger/gat). GAT determines the significance of overlap between genomic annotations though re-sampling within a genomic workspace defined as the mm10 genome, excluding ENCODE blacklist regions and regions of low mappability (ENCODE accession: ENCFF547MET).

### Enrichment of hearing difficulty heritability in open chromatin regions

Enrichment of hearing difficulty heritability in tissue-specific open chromatin regions from our cochlear ATAC-seq experiments, as well as from ENCODE DNase-seq experiments, was examined using stratified LDSC. 1000 Genomes Phase 3 baseline model LD scores (non tissue-specific annotations) described by Finucane, Bulik-Sullivan et al. (2015)^32^ were downloaded from http://data.broadinstitute.org/alkesgroup/LDSCORE/. Open chromatin regions from DNase-seq of mouse tissues and cell types were downloaded from encodeportal.org; accession identifiers for the specific files are shown in Table S14. Regions of the mouse genome identified in these cochlear and non-cochlear open chromatin experiments were mapped to the human genome with the UCSC Genome Browser liftOver tool, using the mm10toHg19 UCSC chain file, requiring a minimum of 50% of base pairs identical between the two genomes.

### Statistical fine mapping and functional annotation of GWAS risk loci

Fine-mapping was performed using a combination of standard annotations and analyses performed with the SNP2GENE function in FUMA v1.3.1 (http://fuma.ctglab.nl)^62^, as well as additional cochlea-specific annotations downstream data integration, as described below. The architecture of risk loci was determined based on LD structure in the 1000 Genomes Phase 3 European-American sample, calculated with PLINK. LD-independent lead SNPs at each locus had p-values < 5e-8. We defined risk loci using a minimum pairwise r2 > 0.6 between lead SNPs and other SNPs. In addition, we set a minimum minor allele frequency of 0.01, and the maximum distance between LD blocks to merge into interval was 250. Subsequently, we selected 613 SNPs for deeper annotation, using a pairwise r^2^ threshold > 0.9 with an LD-independent lead SNP.

Annotations of protein-coding variants were performed in FUMA, using ANNOVAR^65^ with ENSEMBL v92 gene models. Deleteriousness of variants was predicted using CADD v1.3^66^, and we selected variants with a CADD Phred score threshold >= 10.

Regulatory functions were predicted for non-coding variants based on overlap with open chromatin in mouse cochlear epithelial and non-epithelial cells, followed by prediction of target genes based on proximity to transcription start sites (TSS) and chromatin interactions from Hi-C experiments. First, we selected a subset of the 613 risk-associated SNPs that were located +/−500bp of regions homologous to open chromatin in mouse cochlear epithelial and non-epithelial cells. SNPs were annotated to proximal target genes if they were located within 20kb of the TSS from an ENSEMBL v92 gene model.

SNPs were annotated to distal target genes based on chromatin interactions from Hi-C of 20 human tissues and cell types^34^. Hi-C data were processed with FUMA, using Fit-Hi-C^67^ to compute the significance of interactions between 40kb chromosomal segments. Using these data, we identified chromosomal interactions that connect the genomic segment containing each risk-associated, open chromatin-overlapping SNP to distal chromosomal segments. We annotated genes whose transcription start sites were located within these distal segments. We considered chromosomal loops identified in each of the 20 tissues and cell types. As chromosomal contacts differ from tissue to tissue and Hi-C data are inherently noisy, aggregating loops from multiple tissues can lead to false positive signals. To mitigate this risk, we selected a strict p-value threshold for the significance of loops, p < 1e-25, manually determined by inspection of the data to capture one or a few of the strongest loops at each locus.

### Single cell RNA-seq data analysis

Genomic alignment, de-multiplexing, and mapping of unique molecular identifiers (UMIs) mapping to each gene was performed using cellranger. Downstream analyses were performed with the Seurat R package^51^. We filtered cells with <50 or >20,000 UMIs and with >20% of UMIs coming from mitochondrial genes. Counts of UMIs per gene were log normalized. Highly variable genes were identified using the FindVariableGenes function with the following parameters: dispersion formula = LogVMR, minimum = 0.0123, maximum = 3, y cutoff = 0.5. Counts from variable genes were then scaled. We regressed out effects of cell cycle, percent of mitochondrial genes, and number of unique molecular identifiers. The list of cell cycle genes was obtained from the Seurat website: https://satijalab.org/seurat/cell_cycle_vignette.html. We constructed a shared-nearest neighbors graph based on the first 10 principal components of variation in the scaled and normalized expression patterns of variable genes. Cell clusters were identified from the nearest-neighbors group based on Louvain modularity, using the FindClusters() function, with a resolution of 0.6. Clusters were visualized by t-distributed stochastic neighbor embedding (t-SNE) of the first 10 principal components and annotated to known cochlear cell types based on the expression of all marker genes. Cell type specificity for the 50 risk genes was calculated using FindAllMarkers() function, using Wilcoxon tests to compare counts in each cell type to counts of all other cell types in aggregate.

### Meta-analysis of hair cell-specific gene expression for hearing difficulty risk genes

Processed data from GSE60019, GSE71982, and GSE116703 were downloaded and imported into R using the GEOquery R package. log-transformed transcripts per million were fit to linear models, using the lmFit, contrasts.fit, and eBayes functions in the limma R package. The main effect of cell type (hair cells versus all other cells) was calculated in each dataset, separately, controlling for covariates due to age. We then computed a combined meta-analytic p-value for each gene across the three datasets, using Stouffer’s z-score method with equal weights.

### Data Availability

mRNA-seq, ATAC-seq, and single-cell RNA-seq data have deposited in the Gene Expression Omnibus, accession GSE126129.

