## Supplementary material for "Biological insights from multi-omic analysis of 31 genomic risk loci for adult hearing difficulty": Suppl. Tables 1-21, Suppl. Figs. 1-3: Supplementary Figs Tables S3-S6.docx

**Supplementary Figures and Tables**

Figure S1: Q-Q plots of the 4 hearing-related traits with statistically significant heritability.

Figure S2**:** Marker genes used to identify cell types in postnatal day 2 mouse cochlea.

Figure S3: Locus zoom plot of risk-associated SNPs at the chr22q13.1 risk locus reveals LD-independent protein-coding variants in TRIOBP and BAIAP2L2.

Table S1: Heritability of hearing-related traits. (Excel file)

Table S2: Genetic correlations between hearing related traits and non-hearing related traits in UK BioBank.(Excel file)

Table S3: Genomic risk loci for hearing difficulty.

Table S4: Genomic risk loci for background noise problems.

Table S5: Genomic risk loci for hearing aid use.

Table S6: Genomic risk loci for tinnitus.

Table S7. Replication in the UK Biobank for risk-associated SNPs from previous GWAS of hearing difficulty.

Table S8: MAGMA gene-based p-values. (Excel file)

Table S9: MAGMA gene set enrichments for Mendelian deafness genes and genes expressed in cochlear cell types. (Excel file)

Table S10: MAGMA gene set enrichments for genes expressed in tissues from GTEx.

Table S11: GO Term enrichment of hearing difficulty risk loci. (Excel file)

Table S12. Open chromatin regions in cochlear epithelial cells. (Excel file)

Table S13. Open chromatin regions in cochlear non-epithelial cells. (Excel file)

Table S14. Enrichments of hearing difficulty risk in open chromatin regions from cochlear and non-cochlear cell types. (Excel file)

Table S15. Functional annotations of 613 SNPs in strong LD with LD-independent genome-wide significant SNPs at hearing difficulty risk loci. (Excel file)

Table S16: Chromatin Interactions with hearing difficulty SNPs. (Excel file)

Table S17. List of likely causal genes from integrating coding and non-coding functional annotations. (Excel file).

Table S18. Annotation of 50 fine-mapped risk genes at hearing difficulty risk loci. (Excel file)

Table S19: Marker Genes for cochlear cell types in P1 scRNA-seq data. (Excel file)

Table S20. Cell type-specificity analysis for hearing difficulty risk genes. (Excel file)

Table S21: Meta-analysis of risk gene expression in hair cells vs. other cochlear cell types. (Excel file)

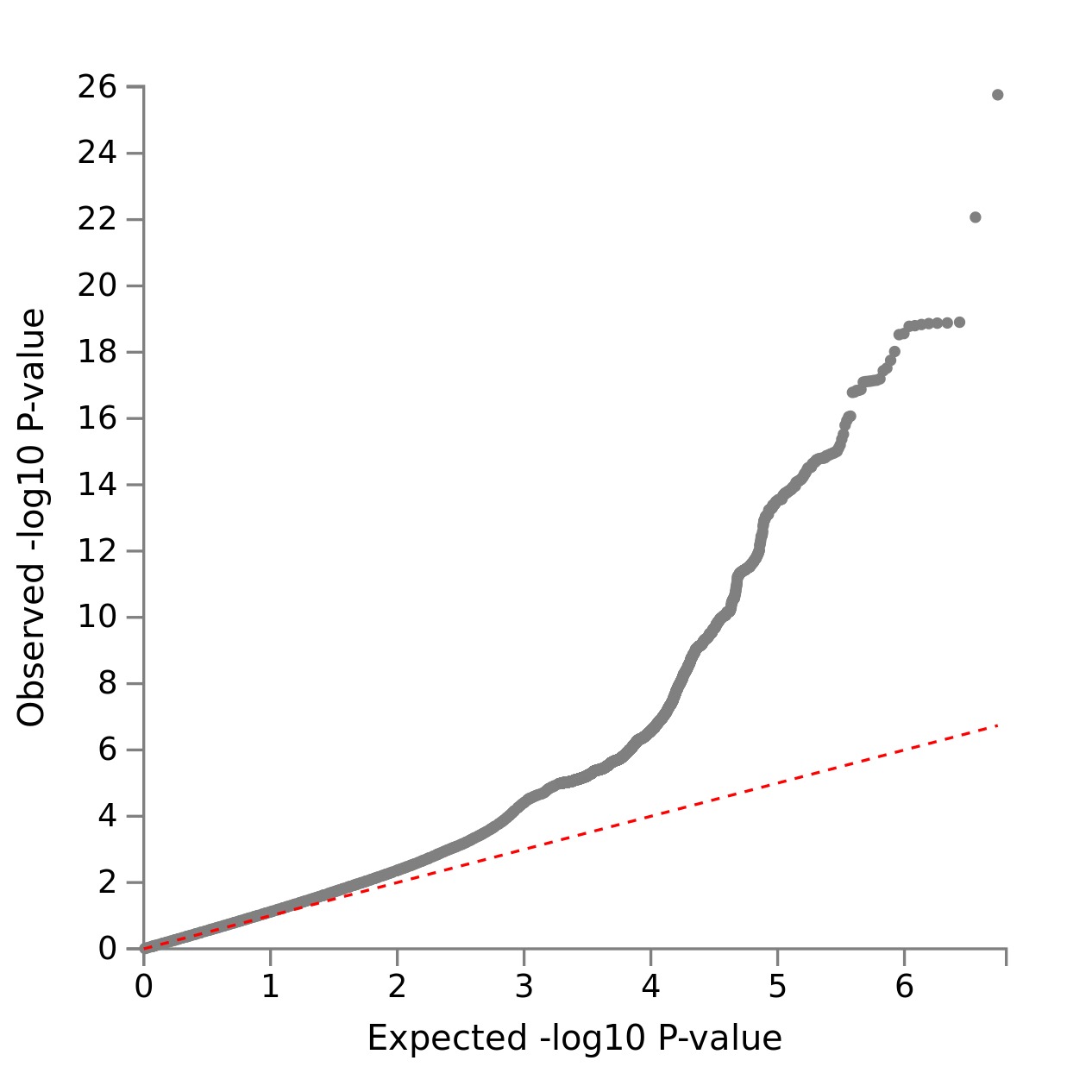

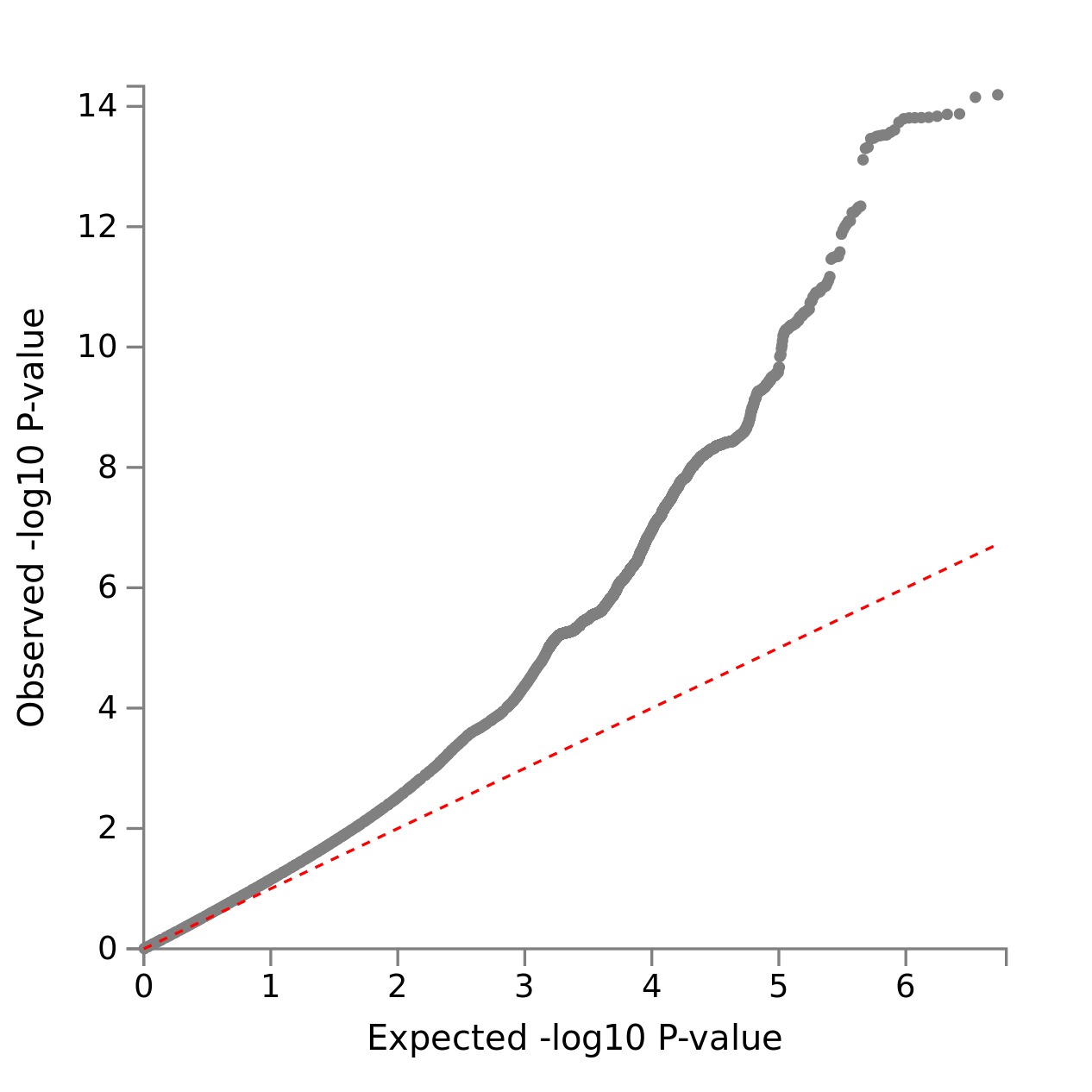

A

B

C

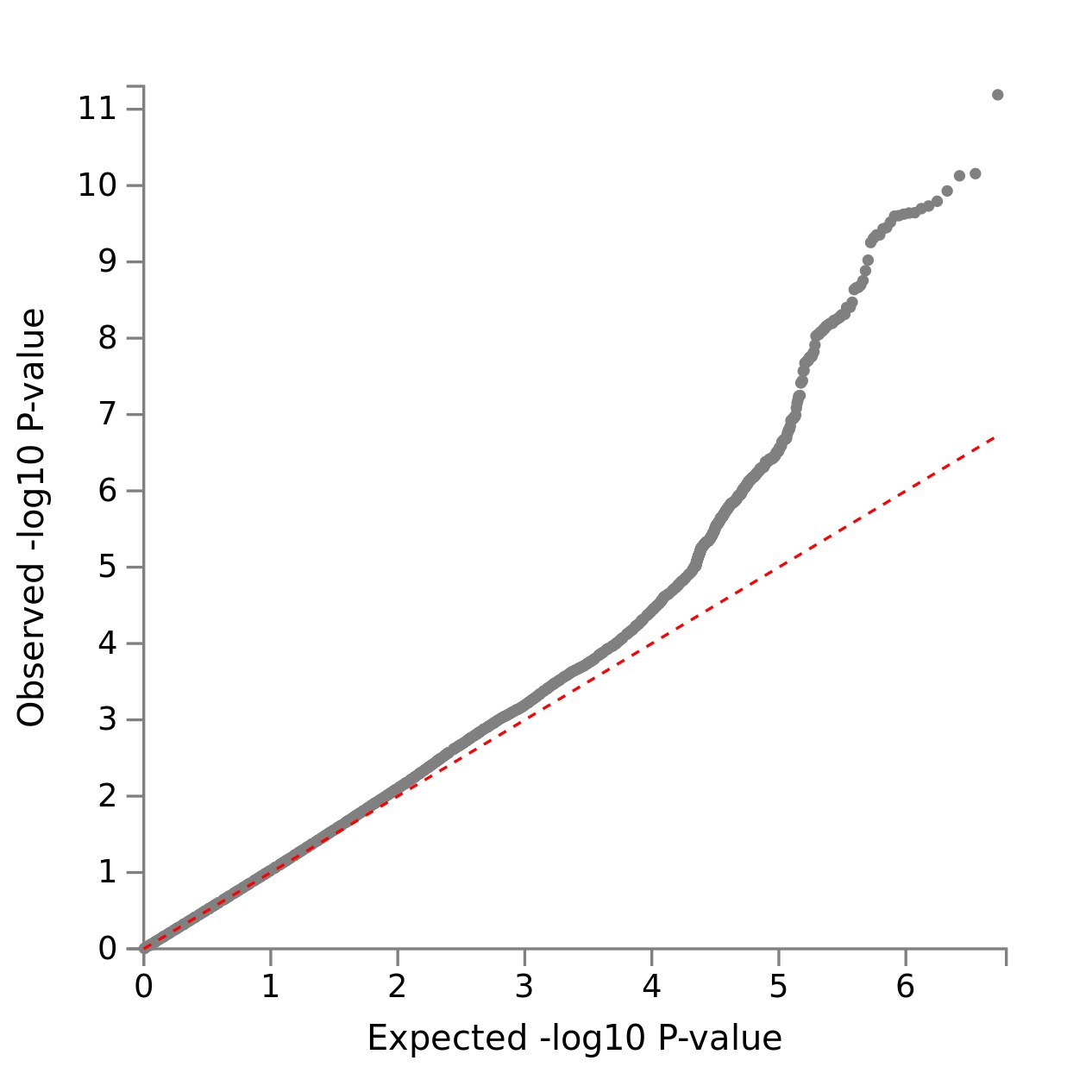

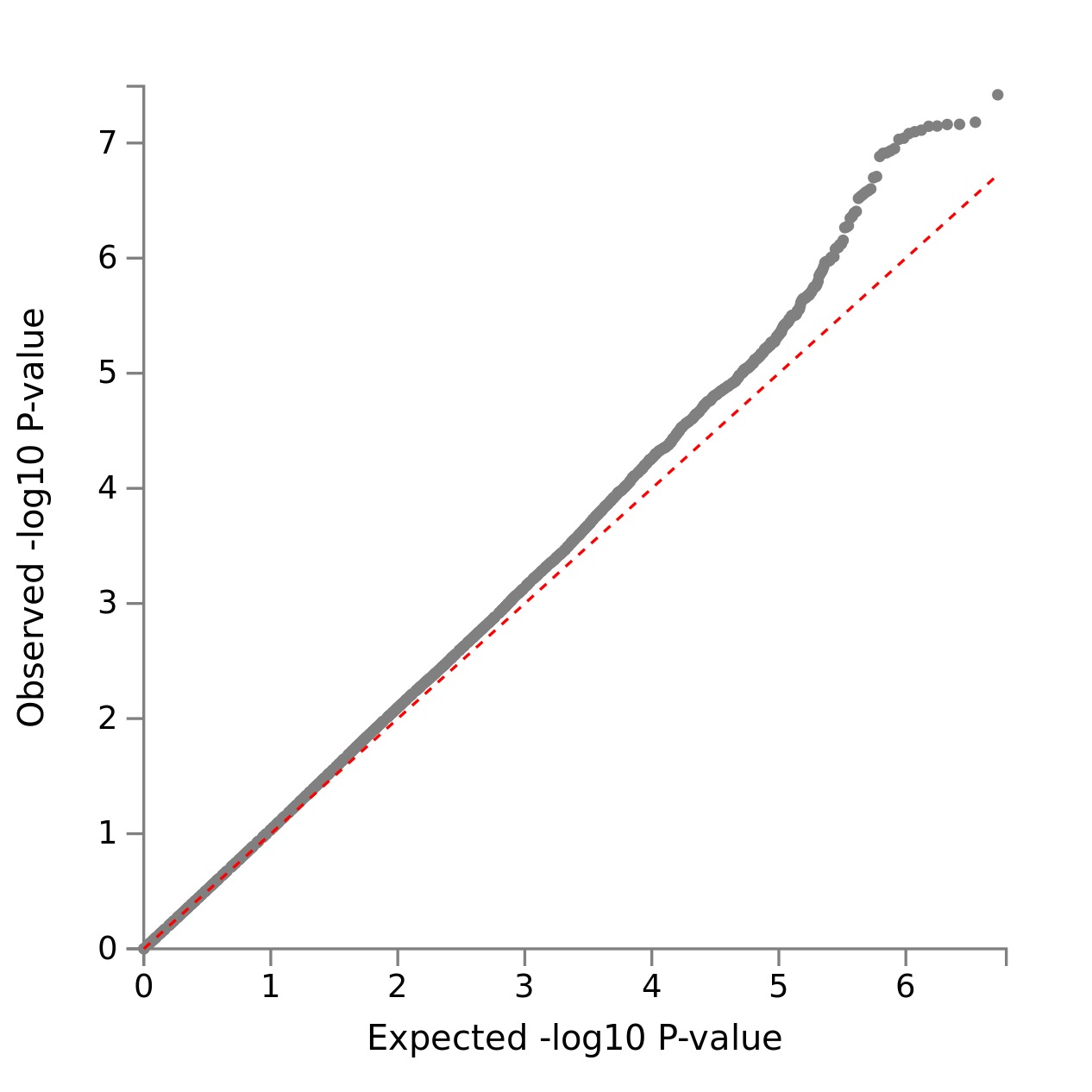

D

**Figure S1**. Q-Q plots of the 4 hearing-related traits with statistically significant heritability. A. 2247_1: “Hearing difficulty/problems: yes” B. 2257: “Hearing difficulty/problems with background noise” C. 3393: “Hearing aid user” D. 4803_11: “Tinnitus: Yes, now most or all of the time.”

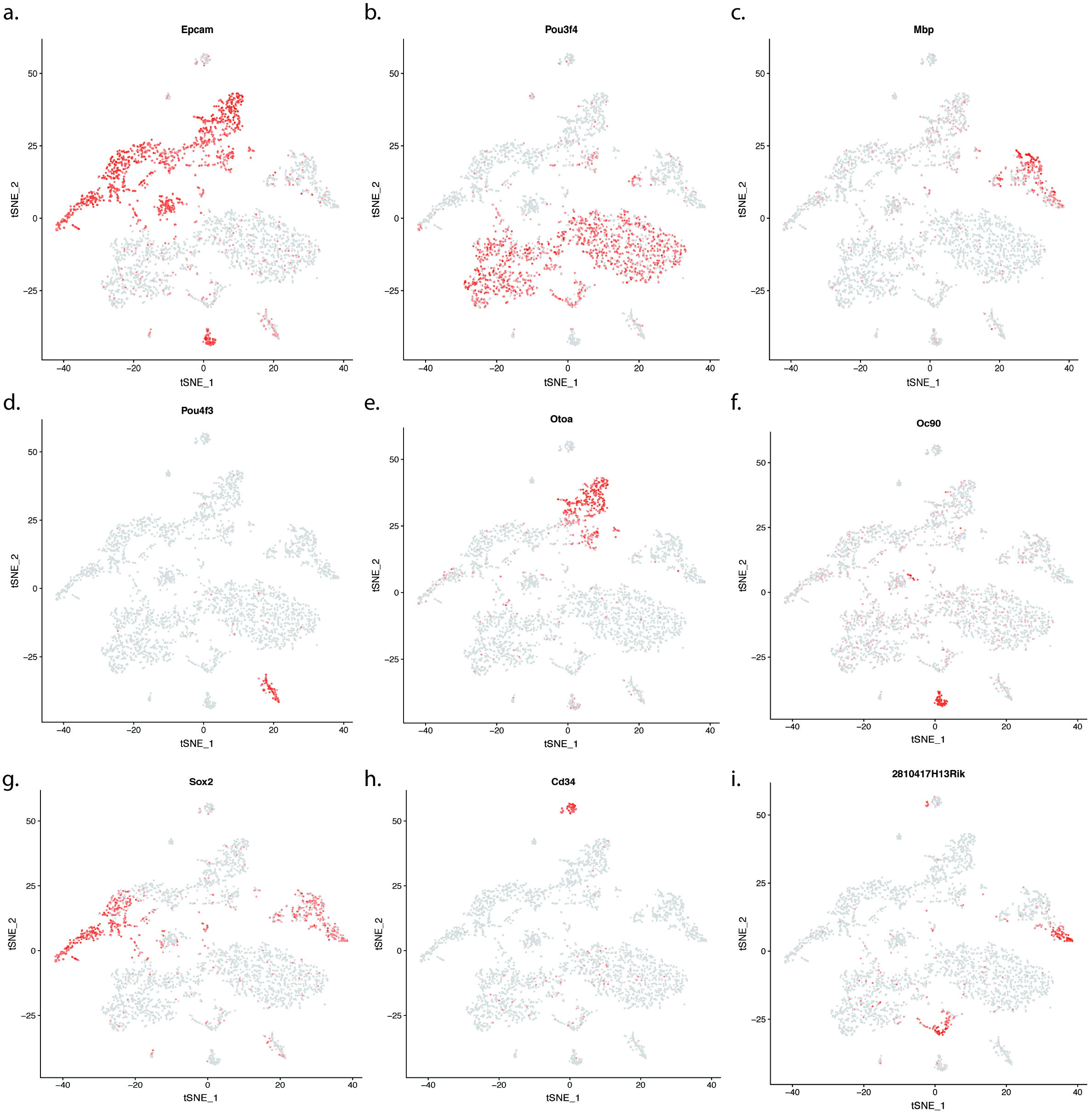

**Figure S2.** Marker genes used to identify cell types in postnatal day 2 mouse cochlea.

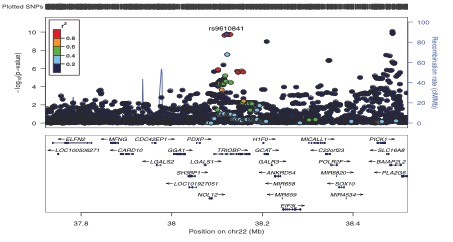

**Figure S3.** Haplotype analysis and functional annotation of risk-associated SNPs at the chr22q13.1 risk locus reveals LD-independent protein-coding variants in TRIOBP and BAIAP2L2.

Table S3. Genomic risk loci for hearing difficulty.

| Locus | uniqID | rsID | chr | pos | p | start | end | Nearest Gene | nSNPs | nGWASSNPs |
| --- | --- | --- | --- | --- | --- | --- | --- | --- | --- | --- |
| 1 | 1:165109131:C:T | rs7525101 | 1 | 165109131 | 1.61E-10 | 165086665 | 165112281 | LMX1A | 22 | 20 |
| 2 | 2:54862003:A:G | rs2941580 | 2 | 54862003 | 1.53E-08 | 54728276 | 54948321 | SPTBN1 | 22 | 17 |
| 3 | 2:208087139:C:T | rs741475 | 2 | 208087139 | 7.72E-09 | 208017033 | 208088987 | KLF7 | 68 | 60 |
| 4 | 3:121712980:C:T | rs3915060 | 3 | 121712980 | 5.84E-09 | 121351315 | 121729262 | ILDR1 | 127 | 109 |
| 5 | 3:182137631:A:C | rs6443802 | 3 | 182137631 | 2.93E-12 | 181935178 | 182210347 | ATP11B | 174 | 140 |
| 6 | 4:17517558:C:T | rs13148153 | 4 | 17517558 | 7.30E-09 | 17517558 | 17530692 | CLRN2 | 5 | 5 |
| 7 | 5:73073075:C:T | rs34929759 | 5 | 73073075 | 1.04E-22 | 72838222 | 73199539 | ARHGEF28 | 382 | 299 |
| 8 | 6:43280713:C:T | rs10948071 | 6 | 43280713 | 9.94E-19 | 43260011 | 43408005 | ZNF318 | 85 | 62 |
| 9 | 6:133789728:A:G | rs9493627 | 6 | 133789728 | 7.66E-10 | 133789728 | 133812872 | EYA4 | 9 | 8 |
| 10 | 6:158505854:C:T | rs6902016 | 6 | 158505854 | 1.04E-12 | 158497717 | 158614172 | SYNJ2 | 157 | 121 |
| 11 | 7:50802434:A:G | rs6968827 | 7 | 50802434 | 7.46E-09 | 50770629 | 50878601 | GRB10 | 56 | 48 |
| 12 | 7:138491839:A:G | rs4732339 | 7 | 138491839 | 2.48E-08 | 138483551 | 138505777 | TMEM213 | 35 | 32 |
| 13 | 8:82657745:A:G | rs74544416 | 8 | 82657745 | 2.25E-08 | 82653644 | 82670771 | CHMP4C | 9 | 8 |
| 14 | 8:141631393:A:G | rs13277721 | 8 | 141631393 | 4.16E-10 | 141604684 | 141704232 | AGO2 | 89 | 63 |
| 15 | 10:73418873:A:G | rs117583072 | 10 | 73418873 | 1.54E-08 | 73418873 | 73418873 | CDH23 | 1 | 1 |
| 16 | 10:94782735:G:T | rs835259 | 10 | 94782735 | 3.78E-08 | 94759464 | 94831523 | EXOC6 | 32 | 26 |
| 17 | 10:126812270:C:T | rs10901863 | 10 | 126812270 | 6.91E-17 | 126783170 | 126812270 | CTBP2 | 3 | 2 |
| 18 | 11:8056913:A:G | rs55635402 | 11 | 8056913 | 1.53E-11 | 8054933 | 8085652 | TUB | 37 | 33 |
| 19 | 11:51422105:A:C | rs61890355 | 11 | 51422105 | 2.39E-08 | 50422724 | 51592021 | OR4A5 | 1331 | 982 |
| 20 | 11:54830428:C:T | rs118176061 | 11 | 54830428 | 3.96E-08 | 54830428 | 55467250 | TRIM48 | 13 | 12 |
| 21 | 11:89017961:A:G | rs1126809 | 11 | 89017961 | 4.94E-15 | 88395396 | 89058101 | TYR | 324 | 260 |
| 22 | 11:118480223:C:T | rs67307131 | 11 | 118480223 | 7.40E-13 | 118478330 | 118735476 | PHLDB1 | 49 | 32 |
| 23 | 14:52514912:C:T | rs1566129 | 14 | 52514912 | 7.15E-11 | 52502345 | 52516601 | NID2 | 39 | 28 |
| 24 | 16:53811788:A:G | rs62033400 | 16 | 53811788 | 9.50E-09 | 53797908 | 53845487 | FTO | 116 | 100 |
| 25 | 16:55492795:A:G | rs78417468 | 16 | 55492795 | 1.33E-08 | 55466943 | 55507988 | MMP2 | 71 | 62 |
| 26 | 17:2574821:A:G | rs12938775 | 17 | 2574821 | 8.10E-09 | 2574821 | 2574821 | PAFAH1B1 | 1 | 1 |
| 27 | 18:44137400:C:T | rs118174674 | 18 | 44137400 | 2.76E-08 | 44137400 | 44137400 | LOXHD1 | 1 | 1 |
| 28 | 18:52636091:C:T | rs4611552 | 18 | 52636091 | 1.92E-08 | 52573169 | 52669732 | CCDC68 | 37 | 32 |
| 29 | 19:2389140:C:T | rs11881070 | 19 | 2389140 | 3.85E-09 | 2367458 | 2394322 | TMPRSS9 | 65 | 51 |
| 30 | 22:38487002:A:G | rs132929 | 22 | 38487002 | 8.04E-11 | 38102301 | 38502927 | BAIAP2L2 | 78 | 62 |
| 31 | 22:50988105:A:G | rs36062310 | 22 | 50988105 | 7.94E-16 | 50973337 | 50988105 | KLHDC7B | 2 | 2 |

Table S4. Genomic risk loci for background noise problems.

| Locus | uniqID | rsID | chr | pos | p | start | end | nSNPs | nGWASSNPs |
| --- | --- | --- | --- | --- | --- | --- | --- | --- | --- |
| 1 | 1:205720483:A:G | rs823116 | 1 | 205720483 | 1.64E-08 | 205646278 | 205799987 | 60 | 49 |
| 2 | 2:208059640:A:C | rs2360675 | 2 | 208059640 | 1.73E-08 | 208031167 | 208088987 | 43 | 36 |
| 3 | 3:124142585:A:G | rs4274696 | 3 | 124142585 | 4.25E-08 | 124142585 | 124186871 | 3 | 3 |
| 4 | 3:182137631:A:C | rs6443802 | 3 | 182137631 | 1.80E-10 | 181938437 | 182150471 | 146 | 115 |
| 5 | 4:17521797:A:G | rs34859220 | 4 | 17521797 | 4.93E-08 | 17517558 | 17530692 | 5 | 5 |
| 6 | 5:73073075:C:T | rs34929759 | 5 | 73073075 | 4.31E-16 | 72839024 | 73198649 | 224 | 169 |
| 7 | 5:174860411:C:T | rs265971 | 5 | 174860411 | 1.46E-08 | 174857623 | 174860411 | 2 | 2 |
| 8 | 6:27198343:G:T | rs67457459 | 6 | 27198343 | 8.88E-09 | 26408472 | 29311921 | 506 | 410 |
| 9 | 6:43342591:A:G | rs4714678 | 6 | 43342591 | 2.23E-17 | 43258464 | 43404511 | 72 | 50 |
| 10 | 6:158513586:G:T | rs2477809 | 6 | 158513586 | 1.81E-08 | 158497717 | 158599382 | 118 | 88 |
| 11 | 7:86237581:G:T | rs1405876 | 7 | 86237581 | 2.65E-10 | 86198818 | 86340497 | 90 | 80 |
| 12 | 8:141629397:A:G | rs13261055 | 8 | 141629397 | 4.16E-09 | 141604684 | 141704232 | 90 | 63 |
| 13 | 10:126812270:C:T | rs10901863 | 10 | 126812270 | 4.64E-11 | 126812270 | 126812270 | 1 | 1 |
| 14 | 11:89037936:C:T | rs1806319 | 11 | 89037936 | 2.12E-10 | 88791508 | 89058101 | 228 | 168 |
| 15 | 11:118478330:A:G | rs7125115 | 11 | 118478330 | 8.27E-09 | 118478330 | 118593218 | 24 | 12 |
| 16 | 12:6871709:A:G | rs11064373 | 12 | 6871709 | 2.83E-08 | 6868482 | 6877721 | 16 | 12 |
| 17 | 17:2574821:A:G | rs12938775 | 17 | 2574821 | 2.19E-12 | 2468731 | 2613762 | 39 | 30 |
| 18 | 19:658335:C:T | rs116941761 | 19 | 658335 | 4.87E-08 | 658335 | 734421 | 10 | 9 |
| 19 | 22:38209650:A:G | rs56265312 | 22 | 38209650 | 4.69E-08 | 38102301 | 38274632 | 10 | 8 |
| 20 | 22:50988105:A:G | rs36062310 | 22 | 50988105 | 9.53E-09 | 50988105 | 50988105 | 1 | 1 |

Table S5. Genomic risk loci for hearing aid use.

| Locus | uniqID | rsID | chr | pos | p | start | end | nSNPs | nGWASSNPs |
| --- | --- | --- | --- | --- | --- | --- | --- | --- | --- |
| 1 | 1:165109131:C:T | rs7525101 | 1 | 165109131 | 5.76E-10 | 165086665 | 165169159 | 63 | 55 |
| 2 | 2:54862003:A:G | rs2941580 | 2 | 54862003 | 2.06E-11 | 54728276 | 54966407 | 176 | 148 |
| 3 | 3:121712980:C:T | rs3915060 | 3 | 121712980 | 1.29E-08 | 121351315 | 121729262 | 127 | 109 |
| 4 | 3:182003490:G:T | rs72622588 | 3 | 182003490 | 5.34E-10 | 181982899 | 182150471 | 133 | 104 |
| 5 | 5:73076024:C:T | rs6871548 | 5 | 73076024 | 9.69E-22 | 72826973 | 73199539 | 481 | 374 |
| 6 | 6:43280713:C:T | rs10948071 | 6 | 43280713 | 8.14E-14 | 43260011 | 43408005 | 76 | 56 |
| 7 | 6:133789728:A:G | rs9493627 | 6 | 133789728 | 1.21E-13 | 133789728 | 133812872 | 9 | 8 |
| 8 | 6:158505854:C:T | rs6902016 | 6 | 158505854 | 1.10E-12 | 158497717 | 158614172 | 157 | 121 |
| 9 | 7:50853151:C:T | rs11238325 | 7 | 50853151 | 2.73E-08 | 50786663 | 50881363 | 117 | 102 |
| 10 | 7:138491839:A:G | rs4732339 | 7 | 138491839 | 1.22E-08 | 138483551 | 138505777 | 35 | 32 |
| 11 | 8:141631393:A:G | rs13277721 | 8 | 141631393 | 3.97E-08 | 141604684 | 141704232 | 89 | 63 |
| 12 | 10:94782735:G:T | rs835259 | 10 | 94782735 | 1.36E-09 | 94759464 | 94831523 | 32 | 26 |
| 13 | 10:126812270:C:T | rs10901863 | 10 | 126812270 | 4.73E-17 | 126812270 | 126812270 | 1 | 1 |
| 14 | 11:8056913:A:G | rs55635402 | 11 | 8056913 | 1.36E-12 | 8053304 | 8099189 | 28 | 27 |
| 15 | 11:51422105:A:C | rs61890355 | 11 | 51422105 | 4.29E-08 | 50422724 | 51592021 | 1331 | 982 |
| 16 | 11:89035134:C:T | rs11018564 | 11 | 89035134 | 3.68E-14 | 88767496 | 89058101 | 283 | 223 |
| 17 | 11:118480223:C:T | rs67307131 | 11 | 118480223 | 4.56E-13 | 118478330 | 118735476 | 90 | 63 |
| 18 | 14:52514912:C:T | rs1566129 | 14 | 52514912 | 3.71E-14 | 52502345 | 52516601 | 39 | 28 |
| 19 | 15:89236386:G:T | rs12441297 | 15 | 89236386 | 1.92E-08 | 89223152 | 89265679 | 37 | 33 |
| 20 | 16:55492795:A:G | rs78417468 | 16 | 55492795 | 7.52E-10 | 55466943 | 55507988 | 71 | 62 |
| 21 | 18:44137400:C:T | rs118174674 | 18 | 44137400 | 2.09E-08 | 44137400 | 44137400 | 1 | 1 |
| 22 | 18:52629054:A:G | rs72930982 | 18 | 52629054 | 2.03E-09 | 52573169 | 52669732 | 37 | 32 |
| 23 | 19:2378830:A:G | rs4417647 | 19 | 2378830 | 4.52E-08 | 2367458 | 2394322 | 45 | 39 |
| 24 | 22:38122448:C:T | rs739137 | 22 | 38122448 | 1.84E-12 | 38102301 | 38502927 | 78 | 62 |
| 25 | 22:50988105:A:G | rs36062310 | 22 | 50988105 | 8.29E-17 | 50973337 | 51157353 | 6 | 6 |

Table S6. Genomic risk loci for tinnitus.

| Locus | uniqID | rsID | chr | pos | p | start | end | nSNPs | nGWASSNPs |
| --- | --- | --- | --- | --- | --- | --- | --- | --- | --- |
| 1 | 1:165109131:C:T | rs7525101 | 1 | 165109131 | 1.25E-08 | 165086665 | 165112281 | 22 | 20 |
| 2 | 2:54817683:A:G | rs6545432 | 2 | 54817683 | 1.63E-09 | 54686740 | 54948321 | 58 | 42 |
| 3 | 3:182138481:A:T | rs4859223 | 3 | 182138481 | 8.41E-09 | 181982837 | 182150471 | 131 | 103 |
| 4 | 5:73073615:C:T | rs11957938 | 5 | 73073615 | 3.10E-17 | 72839024 | 73199539 | 233 | 183 |
| 5 | 6:43280713:C:T | rs10948071 | 6 | 43280713 | 1.72E-16 | 43260011 | 43408005 | 85 | 62 |
| 6 | 6:84365449:C:T | rs217309 | 6 | 84365449 | 4.08E-08 | 84264202 | 84409255 | 64 | 51 |
| 7 | 6:133789728:A:G | rs9493627 | 6 | 133789728 | 2.42E-08 | 133789728 | 133812872 | 9 | 8 |
| 8 | 6:158505854:C:T | rs6902016 | 6 | 158505854 | 4.45E-09 | 158497717 | 158599382 | 118 | 88 |
| 9 | 8:141629397:A:G | rs13261055 | 8 | 141629397 | 5.39E-09 | 141604684 | 141704232 | 90 | 63 |
| 10 | 10:126812270:C:T | rs10901863 | 10 | 126812270 | 7.10E-13 | 126812270 | 126812270 | 1 | 1 |
| 11 | 11:8056913:A:G | rs55635402 | 11 | 8056913 | 4.78E-10 | 8054933 | 8085652 | 16 | 15 |
| 12 | 11:89037064:A:G | rs1847140 | 11 | 89037064 | 4.25E-12 | 88371036 | 89058101 | 410 | 319 |
| 13 | 11:118480223:C:T | rs67307131 | 11 | 118480223 | 4.38E-11 | 118478330 | 118735476 | 49 | 32 |
| 14 | 14:52514912:C:T | rs1566129 | 14 | 52514912 | 3.48E-12 | 52502345 | 52516601 | 39 | 28 |
| 15 | 16:51165891:C:T | rs2111125 | 16 | 51165891 | 2.20E-08 | 51163406 | 51189169 | 17 | 14 |
| 16 | 16:53814363:G:T | rs9972653 | 16 | 53814363 | 3.11E-08 | 53797908 | 53845487 | 116 | 100 |
| 17 | 16:55492795:A:G | rs78417468 | 16 | 55492795 | 2.39E-08 | 55466943 | 55507988 | 71 | 62 |
| 18 | 18:44137400:C:T | rs118174674 | 18 | 44137400 | 2.76E-08 | 44137400 | 44137400 | 1 | 1 |
| 19 | 22:38122448:C:T | rs739137 | 22 | 38122448 | 1.17E-09 | 38102301 | 38502927 | 67 | 55 |
| 20 | 22:50988105:A:G | rs36062310 | 22 | 50988105 | 1.05E-10 | 50988105 | 50988105 | 1 | 1 |
